# Multiple congenital malformations arise from somatic mosaicism for constitutively active Pik3ca signaling

**DOI:** 10.1101/2022.04.05.487130

**Authors:** Elise Marechal, Anne Poliard, Kilian Henry, Mathias Moreno, Mathilde Legrix, Nicolas Macagno, Grégoire Mondielli, Teddy Fauquier, Anne Barlier, Heather C. Etchevers

## Abstract

Recurrent missense mutations of the *PIK3CA* oncogene are among the most frequent drivers of human cancers. These often lead to constitutive activation of its product p110α, a phosphatidylinositol 3-kinase (PI3K) catalytic subunit. In addition to causing a broad range of cancers, the H1047R mutation is also found in affected tissues of a distinct set of congenital tumors and malformations. Collectively termed *PIK3CA*-related disorders (PRDs), these lead to overgrowth of brain, adipose, connective and musculoskeletal tissues and/or blood and lymphatic vessel components. Vascular malformations are frequently observed in PRD, due to cell-autonomous activation of PI3K signaling within endothelial cells. These, like most muscle, connective tissue and bone, are derived from the embryonic mesoderm. However, important organ systems affected in PRDs are neuroectodermal derivatives. To further examine their development, we drove the most common post-zygotic activating mutation of *Pik3ca* in neural crest and related embryonic lineages. Outcomes included macrocephaly, cleft secondary palate and more subtle skull anomalies. Surprisingly, *Pik3ca*-mutant subpopulations of neural crest origin were also associated with widespread cephalic vascular anomalies. Mesectodermal neural crest is a major source of non-endothelial connective tissue in the head, but not the body. To examine the response of vascular connective tissues of the body to constitutive Pik3ca activity during development, we expressed the mutation by way of an Egr2 (Krox20) Cre driver. Lineage tracing led us to observe new lineages that had normally once expressed Krox20 and that may be co-opted in pathogenesis, including vascular pericytes and perimysial fibroblasts. Finally, Schwann cell precursors having transcribed either Krox20 or Sox10 and induced to express constitutively active PI3K were associated with vascular and other tumors. These murine phenotypes may aid discovery of new candidate human PRDs affecting craniofacial and vascular smooth muscle development as well as the reciprocal paracrine signaling mechanisms leading to tissue overgrowth.

## 1 Introduction

Inappropriate activation of signaling components between the cell membrane and its nucleus leads to pathologies with onset at any stage of life, ranging from before birth into old age. These include most cancers, but also numerous, individually rare diseases with a congenital basis.

Nearly two dozen phenotypically disparate overgrowth disorders are mosaic for gene mutations that constitutively activate the phosphatidylinositol 3-kinase (PI3K) pathway (Canaud et al., 2021). A striking majority of these diseases show the same hotspot, activating mutations in the *PIK3CA* gene as over one in eight U.S. cancers (Mendiratta et al., 2021). A large subset of PIK3CA-related disorders (PRDs) is also known as “PIK3CA-related overgrowth syndromes” or PROS, where susceptible tissues such as the cortex (Alcantara et al., 2017), skeletal muscles (Frisk et al., 2019) or facial skin and bone (Couto et al., 2017) develop segmental overgrowth. Because the mutations are usually somatic, the induced overgrowth can be massive and sometimes, life-threatening.

Normally, receptor tyrosine kinase-mediated recruitment and activation of PI3Ks lead to appropriate production of second messengers from lipid substrates. PI3Ks consist of a regulatory p85 subunit and one of three possible 110-kDa catalytic subunits (p110α, p110β, p110δ). *PIK3CA* encodes p110α. Distinct combinations of 3-phosphoinositide substrates produced by PI3Ks and the three AKT protein isoform effectors mediate metabolic, trafficking, growth, survival, proliferation and motility processes to coordinate cellular responses with other signaling pathways. How imbalanced PI3K signaling affects normal homeostasis to cause relatively unchecked cell proliferation before birth, but cancer later in life, is not understood.

In a recent inducible mouse PRD model, penetrance and severity of retinal venous and/or capillary malformations correlated with the process of active angiogenesis (Kobialka et al., 2022). Lesions arose as a function of exposure to normal, age-dependent local growth factor stimuli, rather than of the extent of mosaicism. As seen in human PRD vascular malformations, mosaicism for constitutively active *Pik3ca* entailed the permanent loss of pericytic coverage, while both endothelial proliferation and pericyte loss could be reversed in this model upon administration of an Akt inhibitor (Kobialka et al., 2022). Thus, other symptoms of PRDs may also be temporally constrained in their onset to specific windows of development.

Neural crest (NC) cells are the source of pericytes and vascular smooth muscle in the face, neck, heart and forebrain, including the eyes (Etchevers et al., 2001). They also contribute all corresponding subcutaneous and connective tissues, including facial and frontal bones. Towards the end of the first month of human gestation, NC cells migrate away from the dorsal midline of the future brain and spinal cord to give rise to a wide variety of differentiated cell types (Le Douarin and Kalcheim, 1999). Their dispersion throughout the embryo brings them in contact with changing microenvironments. Errors in NC function lead to a large class of diseases collectively known as neurocristopathies (Bolande, 1974; Etchevers et al., 2019). The normal potential of NC stem cells to both influence surrounding tissues and to differentiate according to context means this population is a prime target for transient growth factor signaling anomalies (Le Lievre and Le Douarin, 1975; Bergwerff et al., 1998; Etchevers et al., 1999, 2001; Müller et al., 2008; Zachariah and Cyster, 2010).

NC specification, migration and differentiation are dependent on PI3K signaling, as shown by loss-of-function studies (Ciarlo et al., 2017; Sittewelle and Monsoro-Burq, 2018). Here, we describe new tools to test the hypothesis that gain-of-function *Pik3ca* mutations in other blood vessel cell lineages than the endothelium itself can also cause vascular malformation syndromes. Our findings both model and extend the PRD spectrum to include effects on the craniofacial skeleton.

## 2 Materials and Methods

### 2.1 Mouse lines

All mice were obtained directly or through Charles River France from the Jackson Laboratories (Bar Harbor, ME, USA) and intentionally outbred over multiple generations to CD-1/Swiss mice purchased from Janvier Laboratories in order to phenocopy human genetic heterogeneity. Knock-in lines included conditional, floxed *Pik3ca*^H1047R^ (RRID:IMSR_JAX:016977) (Adams et al., 2011) or *RdTomato* reporter mice (RRID:IMSR_JAX:007909); transgenics included the *Wnt1-Cre* (RRID:IMSR_JAX:003829), *Krox20-Cre* (RRID:IMSR_JAX:025744) and tamoxifen-inducible *Sox10-CreER^T^* (RRID:IMSR_JAX:027651) lines. 4-hydroxy-tamoxifen (Merck H7904) was solubilized in 100% ethanol, emulsified in sterile corn oil under heated agitation, then the ethanol evaporated in a SpeedVac vacuum concentrator (ThermoFisher) and replaced with corn oil to 10 mg/ mL. 1 mg was administered by intraperitoneal injection (25-35 ng/g body weight). Euthanasia was performed by carbon dioxide inhalation for animals older than 14 postnatal (P) days and by bisection or decapitation for fetuses older than embryonic day 15 and until P14. All mice were housed in individual, ventilated cages with 12-hr light/dark cycles with food and water *ad libitum.*

Mice were genotyped with 50 ng DNA purified from ear punch or tail clips using the primers described in the original reports and Phire Tissue Direct PCR Master Mix (Thermo Scientific) according to manufacturer’s recommendations.

### 2.2 Ethics approval

The animal study was reviewed and approved by the French national animal care and use committee (ACUC) C2EA-14 under the reference 9522-2017040517496865v5.

### 2.3 Histology, immunofluorescence and *in situ* hybridization

Embryos were staged taking embryonic day (E) 0.5 as the morning of the vaginal plug. Tissue biopsies were kept in ice-cold phosphate-buffered saline (PBS) until dissection, fixed in freshly thawed, neutral pH-buffered 4% paraformaldehyde for 20 minutes to overnight depending on tissue size, and rinsed again in PBS. Paraffin blocks were prepared according to standard embedding protocols, sections cut on a Leica microtome at 7 μm, and deparaffinated and rehydrated to PBS through xylene and decreasing ethanol solutions. Alternatively, fluorescent tissues were equilibrated in 15% then 30% sucrose in PBS and positioned in liquid embedding compound (Leica) before snap-freezing in plastic molds over liquid nitrogen. Cryosections were cut at 12 μM onto Superfrost Plus slides, dried, washed in PBS. All immunofluorescent sections were immersed for 20 minutes in 50 mM glycine, 0.1 M ammonium chloride before pre-incubating in a blocking solution of 0.1% Tween-20, 2% fetal calf serum in PBS and diluting the primary antibodies at the indicated concentrations for overnight treatment under Parafilm coverslips at 4°C. Standard procedures were followed for DAPI counterstain, subsequent Alexa Fluor-coupled secondary antibody (ThermoFisher) incubation and mounting with Fluoromount G (SouthernBiotech) under coverslips.

The following primary antibodies were used in this study: rat anti-Pecam1/CD31 (ThermoFisher, RRID:AB_467201), mouse anti-alpha-smooth muscle actin (Sigma-Aldrich, RRID:AB_10979529), rabbit anti-phosphorylated-S6 ribosomal protein (Ser235/236) (Cell Signaling Technologies, RRID:AB_2181035). Standard hematoxylin-eosin (HE; with or without Alcian blue to detect sulfated glycosaminoglycans) staining protocols were followed for designated sections.

### 2.4 Microscopy

Gross anatomy was photographed with a Leica MZ6 dissecting microscope and images captured with a DFC450 camera before analysis using the open-source ImageJ software (v1.53). Histology slides were photographed on a Zeiss AxioScan 7 and immunofluorescent sections on a Zeiss AxioZoom, Apotome or LSM800-Airyscan microscope equipped with Zen software (v2.3 or 3.0). Centroid size of crania was quantified in ImageJ by delimiting the shape above a virtual line from the upper jaw to earlobe to occiput in the sagittal plane (Pilatti and Astúa, 2017).

### 2.5 Micro-X-ray computed tomography (μCT) examination

Late fetal (embryonic day [E]15.5-E20) specimens were genotyped and heads fixed overnight in 4% buffered paraformaldehyde at 4°C. They were stored in PBS + 0.1% w/v sodium azide before imaging on a X-ray micro-CT device (Quantum FX Caliper, Life Sciences, Perkin Elmer, Waltham, MA) hosted by the PIV Platform, EA2496, Montrouge, France. The X-ray source was set at 90 kV and 160 μA. Tridimensional images were acquired with an isotropic voxel size of 20 μm. Non-mineralized tissues were visualized after impregnating with Lugol’s solution.

Tiff image stills were extracted from Dicom data frames using licensed 64-bit Irfanview imaging freeware (v4.59, http://www.irfanview.com/). 2D measurements were made in ImageJ after manual segmentation using contrast thresholding.

### 2.6 Statistical methods

Animals of a given genotype from three or more litters were used for all observations. Groups were compared for significant differences between the means of small samples drawn from normally distributed populations using Student’s t-test and plots generated with Prism version 9.4.1 (GraphPad).

## 3 Results

### 3.1 Constitutively active Pik3ca in most neural crest leads to perinatal death and craniofacial malformations

In order to understand the effects of PI3K signaling in neural crest (NC) derivatives, we mated conditional, floxed *Pik3ca*^H1047R^ and/or Tomato *(RdT)* reporter (Madisen et al., 2010) knock-in lines with *Wnt1-Cre* transgenic mice, which express Cre recombinase in nearly all NC-derived cells from pre-migratory stages onwards, as well as along the dorsal midline of the central nervous system (CNS), ventral forebrain and most midbrain neuroepithelium (Echelard et al., 1994; Danielian et al., 1998). This cross induced somatic mosaicism for constitutively active PI3K in only those animals carrying both a floxed *Pik3ca* and a *Cre* allele. Mosaicism was restricted to tissues having expressed the recombinase at any point before phenotypic analysis (Sauer, 1998).

No *Wnt1-Cre*; *Pik3ca*^H1047R/+^ mice were recovered at weaning; in fact, all *Wnt1-Cre; Pik3ca*^H1047R/+^ mice died within the first day after birth (n=12 mutants and 58 live sibs). The proportion of mutants at birth was not statistically different than expected from Mendelian ratios (X^2^[1,140] = 1.299, p = 0.254). The dead neonates had no visible belly milk spot and notably large heads; three had cleft palates (**Figure 1**). In order to better understood the causes and onset of death, we harvested mice at unaffected control littermates expressing only one or neither of the *Wnt1-Cre* or *Pik3ca*^H1047R/+^ alleles.

**Figure 1.**
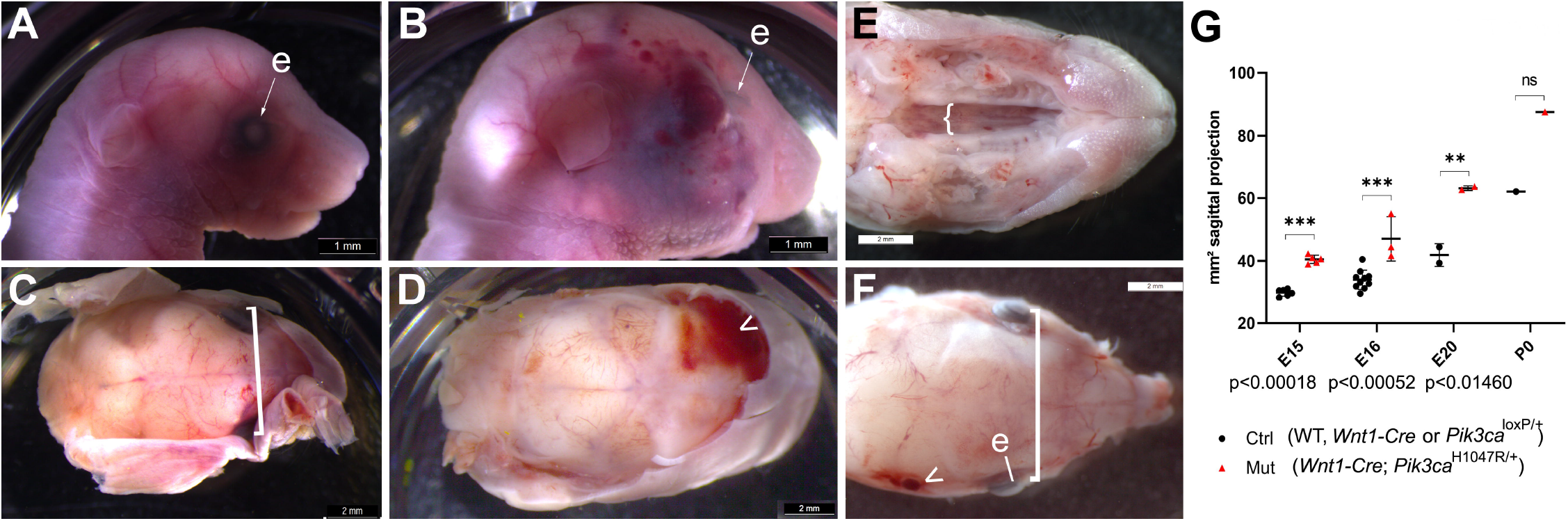
Mutants compared to their unaffected control littermates expressing only one or neither of the *Wnt1-Cre* or *Pik3ca*^H1047R/+^ alleles. (**A**) At E20, fetal head from control littermate of (**B**) at same magnification, facing right. (**B**) At E20, mutant fetuses and newborns had visible megalencephaly, displaced eyes above an enlarged maxillary primordium and skull vault, larger external ears, a longer nose, and facial vascular malformations or tumors, sometimes hemorrhagic (arrowheads). Bar, 1 mm. (**C**) Distinct E20 control fetus, skin removed and relative interocular distance indicated in square bracket. Bar, 2 mm. (**D**) Distinct E20 mutant fetus, skin removed to see hemorrhage from ruptured vascular lesions. Bar, 2 mm. (**E**) *Wnt1-Cre^+/o^*; *Pik3ca*^H1047R/+^ mutant newborn at P0, view of cleft palate (bracket) after removal of skin and jaw. Bar, 2 mm. (**F**) *Wnt1-Cre*^+/o^; *Pik3ca*^H1047R/+^ mutant newborn at P0, coronal view of cleft palate (bracket) after removal of skin and jaw, relative interocular distance indicated in square bracket at same scale as (**C**). Arrowhead, vascular lesion. Bar, 2 mm. (**G**) Maximum sagittal plane surface projection of lateral photographs from embryos at the indicated stages between embryonic day (E)15 and birth (P0), of either control (ctrl) littermate or *Wnt1-Cre^+/o^*; *Pik3ca*^H1047R/+^ (mut) genotype. Through the last third of gestation, mutant heads were significantly larger than controls, as was the head of the sole neonate recovered. ctrl, control; e, eye; mut, mutant; ns, not significant.

From E15.5 onwards, mutant crania were all visibly larger than controls (**Figure 1G**, n=12 mutants, 30 controls; p<0.01 at E20 and <0.001 at E15.5-E16; **Figure 2**). Body sizes appeared unaltered at all time points, but mutants had varying degrees of facial soft and calcified tissue hyperplasia as seen in micro-computed tomography (micro-CT; **Figure 2**). At E20, just prior to birth, mutant fetuses were macrocephalic with enlarged frontal, nasal as well as maxillary bones, relative to controls. Strikingly, the parietal bones overlying the burgeoning midbrain, of mesodermal origin in mice (Jiang et al., 2002a), were hypocalcified in mutants relative to controls, while adjacent craniofacial skull components of NC origin were more calcified than usual at this stage and hyperplasic (**Figure 2A-F**).

**Figure 2.**
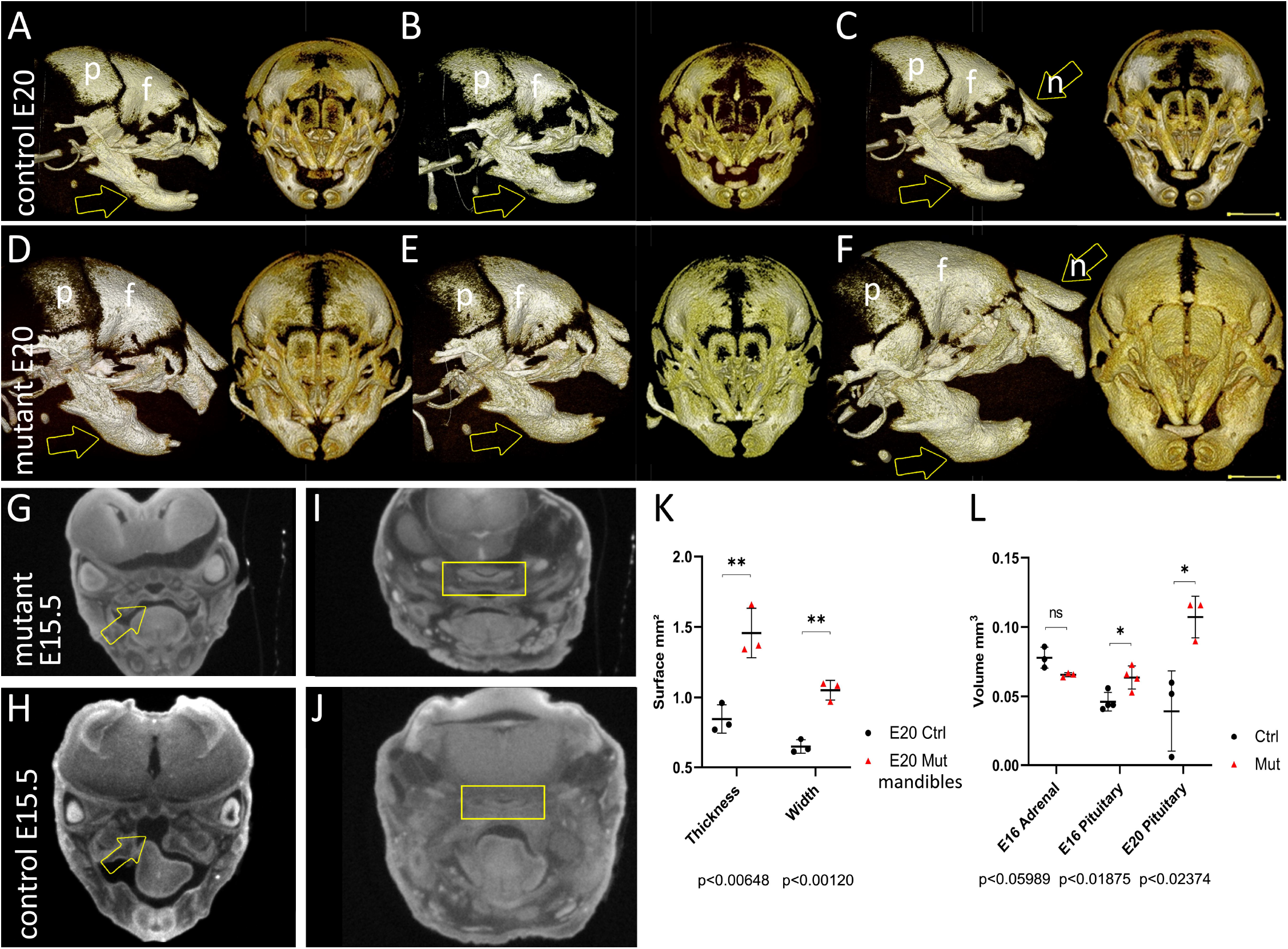
Micro-computed tomography (micro-CT) of skull development in *Wnt1-Cre^+/o^*; *Pik3ca*^H1047R/+^ mice at late fetal stages. (**A-C**) 3D projections in frontal and sagittal views of the more advanced mineralization of three representative skulls of control littermate fetuses at E20.5 relative to mutants in (**D-F**), within neural crest-derived craniofacial bones. Arrows indicate point of measurements of mandibular bone thickness in (**K**). (**G**) Control littermate fetus at E15.5 relative to mutant in (**H**) –frontal micro-CT section through eyes, brain, palate (arrow), tongue and jaw. (**H**) Mutant littermate embryo of E15.5 embryo in (**G**)–frontal micro-CT section through eyes, brain, cleft palate with malpositioned tongue (arrow) and jaw. (**I**, **J**) Frontal sections through the pituitary gland (yellow rectangle) in a control (**I**) or *Wnt1-Cre^+/°^*; *Pik3ca*^H1047R/+^ (**J**) fetus at E15.5. (**K**) Measurements from the ramus base to just anterior to the molar alveolus, in both rostrocaudal thickness (height) and mediolateral width, show significant enlargement of mutant mandibles by birth. (**L**) Segmentation analysis of the volumes of the adrenal and pituitary glands at E16 and the pituitary at E20, showing that unlike the adrenal gland, there is a significant increase in pituitary size at the end of gestation. f, frontal; n, nasal; p, parietal; ns, not significant; * p<0.05; ** p<0.01.

Measurements taken in three *Wnt1-Cre*; *Pik3ca*^H1047R/+^ fetuses at E20 showed significantly thicker and wider mandibles than in controls (**Figure 2A-F**). Mutant mandibular bodies measured 1.457±0.176 mm high and 1.052±0.070 mm wide from the ramus base to just anterior to the molar alveolus, as compared to three control littermates at 0.847±0.101 mm high and 0.649±0.049 mm wide, respectively (p<0.01 in each dimension, **Figure 2K**).

Although only a quarter of the *Wnt1-Cre*; *Pik3ca*^H1047R/+^ fetuses had a cleft secondary palate between E15.5 and birth, no such clefting was ever observed in controls (p<0.05). Possible additional explanations for perinatal death in mutants included defects in vital organs dependent on neural crest-derived components for their morphogenesis. Among these were the adrenal or pituitary glands, the latter of which control adrenal function through the function of its corticotropic cells. Subtle cardiac malformations were an additional possibility: tricuspid valve insufficiency, leading to thickening of ventricular or septal walls, or patent ductus arteriosus (Yajima et al., 2013). We therefore compared these organ sizes on soft tissue contrast-enhanced micro-CT sections from E15.5-E20 fetuses (n=10 mutants vs 10 controls, **Figure 2L**).

### 3.2 *Wnt1-Cre*; *Pik3ca*^H1047R/+^ mice develop variable ocular and pituitary malformations

No significant differences in cardiac or adrenal sizes in micro-CT frontal 2D cross-section were measured at any age in *Wnt1-Cre*; *Pik3ca*^H1047R/+^ mice relative to control littermates. In contrast, the pituitary gland was variably misshapen and significantly smaller in section at E15.5-E16 (n=5, p<0.01). However, volumetric analysis (**Figure 2L)** showed that by birth, the mutant pituitary gland (E16: 0.064±0.009 mm^3^, n=4; E20: 0.107±0.015 mm^3^, n=3) had grown significantly larger than controls at the same age (0.046±0.007 mm^3^, p<0.05; and 0.040±0.029 mm^3^; p<0.05, respectively).

NC begin to migrate into the head after E8.5 (Kaucka et al., 2016); at E9.5, we observed that initial cranial and trunk-level NC migration appeared to be unimpeded in *Wnt1-Cre*; *Pik3ca*^H1047R/+^; *RdT* mice versus their *Wnt1-Cre*; *RdT* littermates (**Figure 3(A-B)**). Likewise, at E13.5 (**Figure 3(C-H)**), palatal shelves, head size and the position of the tongue had not yet developed obvious morphological differences.

**Figure 3.**
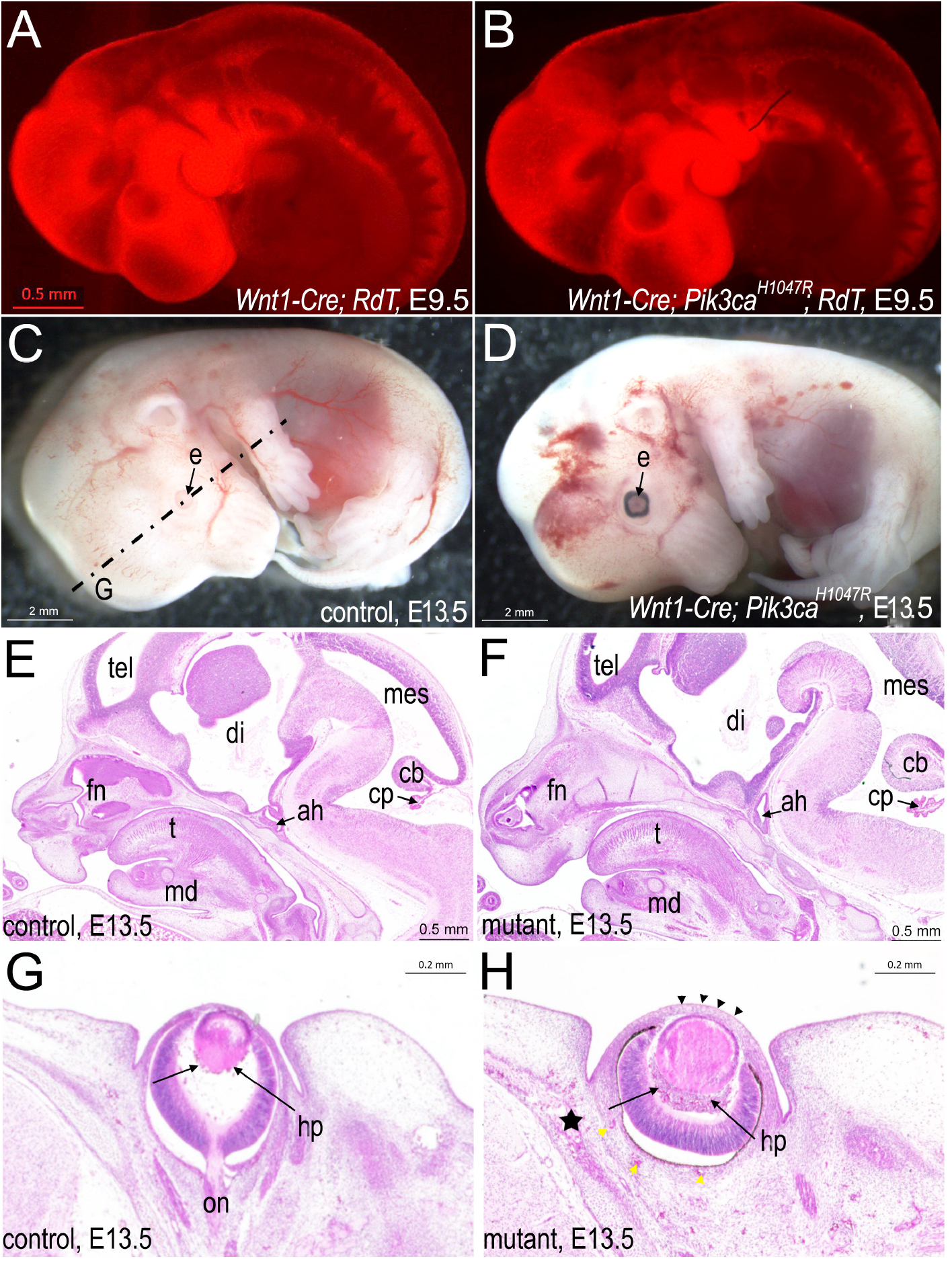
Phenotypes of *Wnt1-Cre^+/°^*; *Pik3ca*^H1047R/+^ embryos from E9.5 to E13.5. (**A**) When lineage-traced by co-expression of a *RdT*^+/o^ allele to transcribe Tomato red fluorescent protein in cells having expressed Wnt1, *Wnt1-Cre*^+/o^; *RdT*^+/o^ embryos at E9.5 show the normal distribution of neural crest (NC) mesenchyme in the face and pharyngeal arches. Bars = 0.5 mm for A, B. (**B**) *Wnt1-Cre^+/o^*; *Pik3ca*^H1047R/+^; *RdT*^+/o^ embryos at E9.5 show unaltered distribution of NC-derived mesenchyme in the pharyngeal arches, frontonasal bud or body. (**C**) Left side of E13.5 control littermate to (**D**). Lack of pigment in retinal pigmented epithelium of eye is normal for a mouse that would have been born albino (Tyr^c^/ Tyr^c^), a background allele (http://www.informatics.jax.org/allele/MGI:1855976). Dotted lines indicate plane of frontal section in G. Bars = 2 mm for C, D. e, eye. (**D**) Left side of E13.5 *Wnt1-Cre^+/o^*; *Pik3ca*^H1047R/+^ littermate to (**C)**, showing vascular anomalies and cerebral hemorrhage. Probable hyaloid plexus malformation visible also in H. (**E**) Paraffin mid-sagittal section of control littermate head of (F) at E13.5, stained with HE. Bars = 0.5 mm for E, F. Abbreviations as in F. (**F**) Paraffin mid-sagittal section of mutant head at E13.5 showing enlarged frontonasal and mandibular (md) tissues, cerebellum (cb) and choroid plexus (cp), as well as a malpositioned adenohypophysis (ah) relative to the infundibulum, stained with HE. di, diencephalon; mes, mesencephalon; t, tongue; tel, telencephalon. (**G**) Paraffin frontal section of control head at E13.5, left eye at level of optic nerve (on), stained with HE. Bar for G, H = 0.2 mm. hp, hyaloid plexus. (**H**) Paraffin frontal section of mutant littermate of (G) at E13.5, left eye, showing lens hypertrophy, hyaloid plexus malformation (arrows), thickened corneal epithelium (black arrowheads) and scleral (yellow arrowheads) and perichondral vascular malformation (star), stained with HE.

However, at E13.5, the eyes already appeared subtly malpositioned relative to other head structures (**Figure 3(C-D)**). In histological section, lens coloboma and microspherophakia, a lack of primary fibers, and a thickened cornea were already evident in mutants (**Figure 3(I-M)**).

### 3.3 PI3K signaling in NC-derived vascular smooth muscle induces venous malformations

One striking and constant feature of *Wnt1-Cre*; *Pik3ca*^H1047R/+^ mutants were their craniofacial vascular malformations, also observable from E13.5 onwards (**Figure 4**). These most resembled venous malformations in that they were non-pulsatile (known as “low-flow” (Olivieri et al., 2016)), circumscribed congenital lesions within the NC-derived dermis over the frontal bones and, frequently, in the maxillary and retroorbital regions (**Figure 4 (A-D)**). The malformations usually contained thromboses and were also observed in the heart, in both ventricles and atria, but not in the trunk.

**Figure 4.**
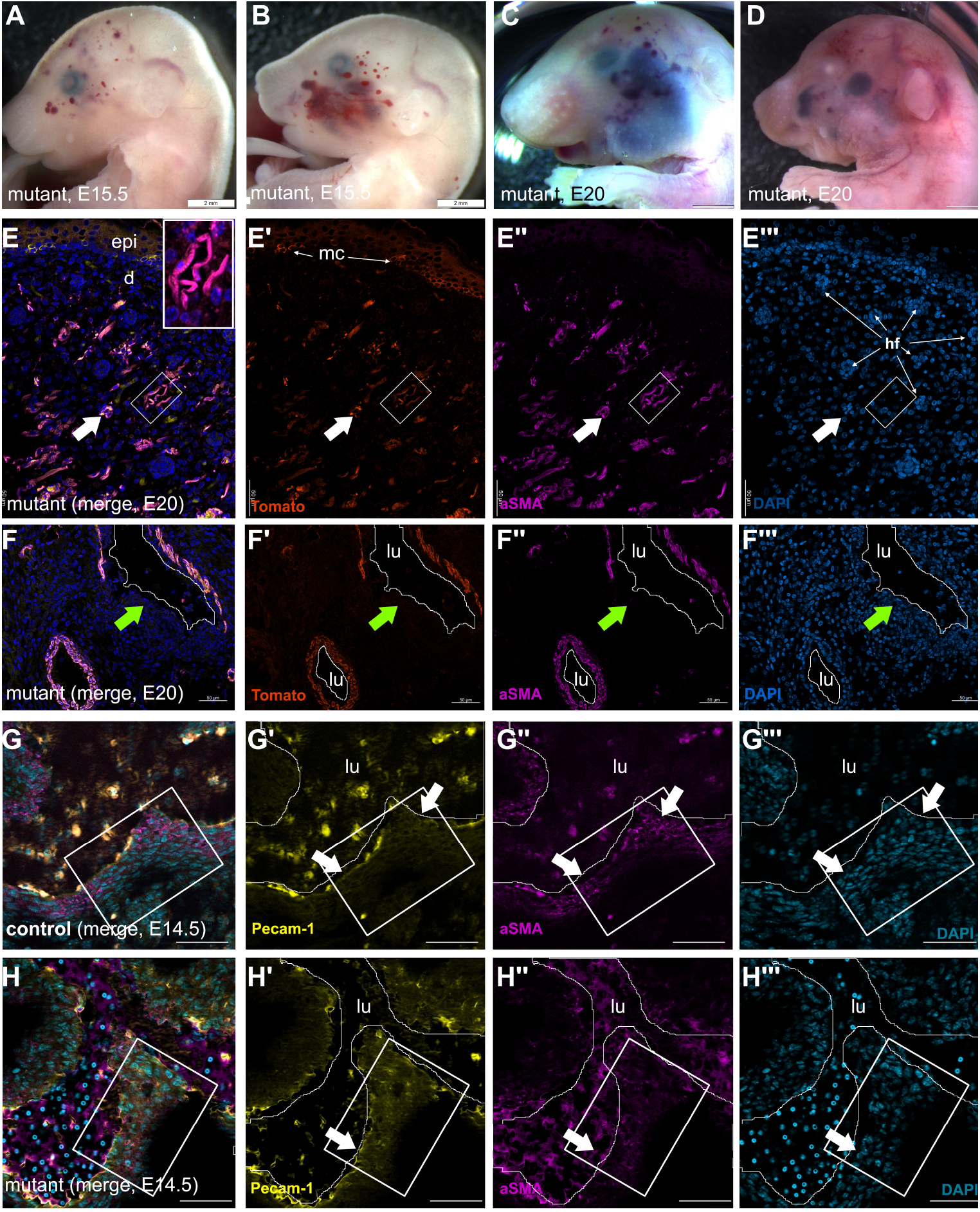
Vascular lesions are present before birth with disorganized smooth muscle or connective elements in *Wnt1-Cre*; *Pik3ca*^H1047R/+^ mutants. (**A**) Mutant embryo at E15.5 with periocular vascular anomalies. (**B**) Mutant littermate of (**A**) with segmental vascular anomalies in the maxillary region. (**C, D**) Mutant E20 fetuses of distinct litters. (**C**) was dead *in utero* and had a vascular lesion on the mandible. All show maxillary and posterior periocular vascular anomalies and megalencephaly. Scale bars A-D, 2 mm. (**E**) Control littermate and (**F**) mutant pulmonary trunk; merged immunofluorescence (**E’**, **F’**): yellow, endothelial marker Pecam-1 (CD31); (**E’’**, **F’’**) purple, smooth muscle marker, α-smooth muscle actin (aSMA); (**E’’’**, **F’’’**) blue, nuclear marker DAPI. Bars E, F = 50 μm. Despite maintenance of endothelium (arrows) in the mutant great artery, boxed areas indicate disruption of the laminar smooth muscle organization of the vascular wall at E14.5. (**G**) Cellular organization around an intracardiac vascular anomaly at E20.5, showing a discontinuous (arrow) smooth muscle layer of mutant NC origin in a distinct litter from the ones represented in C, D. (**H**) Facial skin of E20.5 mutant fetus (littermate of G) with numerous double-labeled, small capillary anomalies in upper dermis (arrow) around hair follicles. (**G’, H’)**: Tomato fluorescent protein; (**G’’, H’’**): smooth muscle marker, α-smooth muscle actin (aSMA); (**G’’’, H’’’**) nuclear marker DAPI in blue. Bars G, H = 50 μm.

In order to examine the composition of these abnormal vascular structures, we examined the distribution of the mature pericyte and smooth muscle marker, α-smooth muscle actin (aSMA) and in *Wnt1-Cre*; *Pik3ca*^H1047R/+^ mutants using immunofluorescence (**Figure 4 (E-H))**. A cutaneous vascular malformation in the skin over the parietal bone in a late fetus at E20.5 (**Figure 4 (E))** showed a profusion of aSMA-expressing cells in abundant dermal blood vessels surrounding the hair follicles (**Figure 4(E’’)**). These also expressed Tomato (**Figure 4(E’)**) and were therefore contributed by the mutation-bearing neural crest cells. Because the omnipresent thromboses were autofluorescent in most channels, we examined other vascular anomalies using immunofluorescence.

Within the late intracardiac lesions, *Wnt1-Cre*; *Pik3ca*^H1047R/+^; *RdT* mutants also showed co-expression of the Tomato NC lineage marker with aSMA in a discontinuous manner around the large vascular lacunae (**Figure 4(F), arrow)**. Smooth muscle cells in the arterial trunk, derived from posterior rhombencephalic NC cells, in E14.5 controls present a concentric, lamellar arrangement of aSMA-expressing cells within the vascular wall (**Figure 4(G)**), **boxed region**), overlying cells expressing the endothelial marker Pecam-1/CD31 (**Figure 4(G’)**). In mutants, where aSMA was present at the base of the arterial trunk, the smooth muscle cells were already partially disorganized and cuboid by E14.5 (**Figure 4(H)**).

These findings indicate that constitutive PI3K activation in neural crest-derived mesenchyme, in both the head and heart where it is usually competent to contribute connective tissues, perturbs their organization and functional roles rather than cellular identity.

### 3.4 PI3K signaling in muscle leads to widespread, progressive vascular anomalies

Despite the striking and lethal phenotype of Pik3ca constitutive activity in mesectodermal cephalic NC, we did not observe any major morphological changes in the trunk in *Wnt1-Cre*; *Pik3ca*^H1047R^ mutants either in whole mount or at birth after autopsy. This included the NC derivatives of the adrenal medulla and the enteric, autonomic or sensory nervous systems.

We wondered if a more subtle cardiac defect in the Wnt1-Cre model, such as valvular insufficiency rather than a major arterial trunk malformation, could be a cause for perinatal death. Neural crest-specific ablation of the conserved zinc finger transcription factor Krox20 (Egr2)-expressing cells has been shown to recapitulate the tricuspid valve hyperplasia and bicuspid aortic valves found in *Krox20* loss-of-function mutants, without affecting the position or separation of the outflow tract itself (Odelin et al., 2018). In order to examine this subset of cardiac NC, we therefore crossed the conditional, floxed *Pik3ca*^H1047R^ and/or Tomato *(RdT)* reporter knock-in lines to the *Krox20-Cre* transgenic line (Voiculescu et al., 2000).

Krox20 has previously been shown to be expressed in connective tissues of both neural crest and non-neural crest origins throughout the head and body, as well as in myelinating Schwann cells throughout life (Voiculescu et al., 2000; Maro et al., 2004). We therefore examined *Krox20-Cre^+/o^*; *Pik3ca*^H1047R/+^ mice for signs of cardiac insufficiency from myxomatous valves, and also for skeletal defects or for peripheral neuropathy.

*Krox20-Cre*; *Pik3ca*^H1047R/+^ mice were initially healthy and viable with no apparent skeletal differences, but over time post-weaning developed palpable lumps under the skin on the back, leg or tail and had soon reached a humane endpoint. Upon autopsy, we discovered widespread, lobular vascular structures filled with coagulated blood in the subcutaneous *panniculus carnosus* muscle, but also in and around skeletal, cardiac and smooth muscles, the lungs, the reproductive organs and many other densely vascularized tissues (**Figure 5** (**A-D, G-T**). These vascular lesions had cavernoma-like fibrous septa, and the adjacent nerves were surrounded by loose fat. No phleboliths were observed and hearts appeared normal. Myelinated sciatic nerves showed no macroscopic or functional differences between mutant and control mice. However, mutant mice all developed striking splenomegaly (**Figure 5**(**E, F**)).

**Figure 5.**
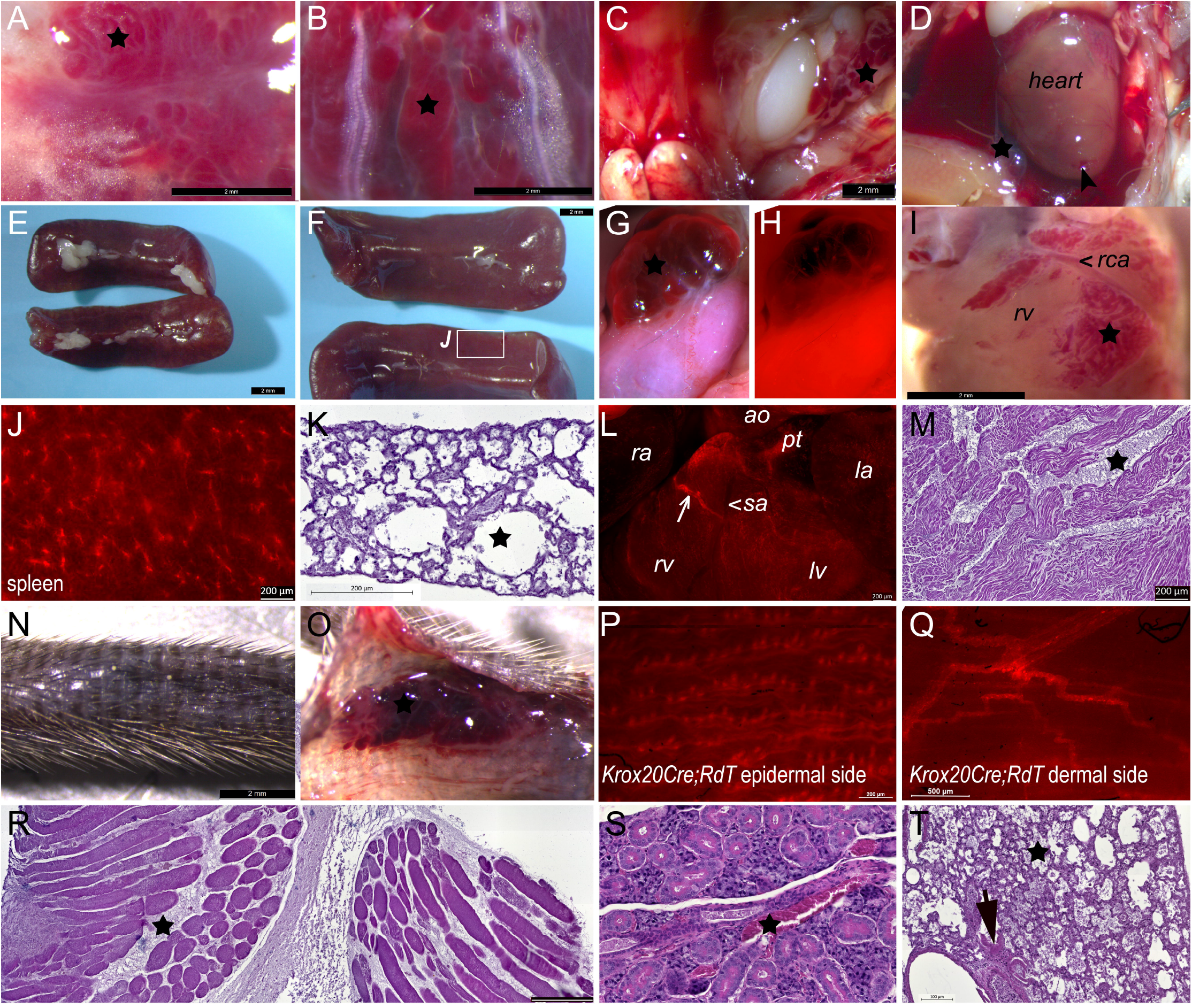
Anatomy and histology of vascular anomalies in *Krox20-Cre^+/o^*; *Pik3ca*^H1047R/+^ adult mutant mice. (**A**) Subcutaneous vascular anomaly (star), epaxial *(longissimus dorsi*) muscle. Bars A-I: 2 mm. (**B**) Subcutaneous vascular anomaly (star), quadriceps. (**C**) Vascular anomalies (star) around left gonad. (**D**) Mutant heart, small vascular anomaly at apex (arrow) and an extracardiac anomaly (star). (**E**) Control littermate spleens of those in (**F**). (**F**) *Krox20-Cre^+/°^*; *Pik3ca*^H1047R/+^; *RdT*^+/o^ spleens, same scale as (E). Boxed region enlarged in (**J**). (**G, H**) Vascular tumor (star) around gonad from different mouse than in (**C**), *Krox20-Cre^+/°^*; *Pik3ca*^H1047R/+^; *RdT^+/o^*. The fluorescent fibroblasts in gonad and lesional septa in (**H)** had expressed Krox20 and thereby, constitutively active Pik3ca. (**I**) Intracardiac vascular anomalies (star) were present in all mutant adults examined. (**J**) *Krox20-Cre^+/°^*; *Pik3ca*^H1047R/+^; *RdT*^+/o^ spleens had numerous fluorescent ramifications consistent with reticular fibers, peripheral nervous or perivascular elements. Bar, 200 μm. (**K**) Mutant femoral bone marrow in an adult mouse that had spontaneously died with multiple vascular anomalies (star) was hypocellular with increased density of vascular sinuses rather than adiposity. Bar, 200 μm. (**L**) Heart from a *Krox20-Cre^+/o^*; *Pik3ca*^H1047R/+^; *RdT^+/o^* mouse showing recombined cells in a fine meshwork throughout the myocardium of all chambers with increased density in the ventral pulmonary trunk and strong expression in the outer wall of a sympathetic nerve (arrow). Bar, 200 μm. (**M**) Typical histology of vascular lacunae (star) in the ventricular wall. Bar, 200 μm. (**N**, **O**) Subcutaneous vascular tumor (star) of tail of *Krox20-Cre^+/o^*; *Pik3ca*^H1047R/+^ mouse. Bar, 2 mm. (**P**) External view of lineage-traced mutant *Krox20-Cre^+/o^*; *Pik3ca*^H1047R/+^ hairy skin. Fluorescent, lineage-traced cells having once expressed Krox20 are visible at the base of each hair follicle and in the vascular pericytes of capillaries underlying them. Bar, 200 μm. (**Q**) Internal view of lineage-traced mutant hairy skin. The *panniculus carnosus* muscle had expressed *Krox20* (striations) and the peripheral nerves express *Krox20Cre*-driven Tomato even more strongly in their myelinating Schwann cells. Bar, 0.5 mm. (**R**) Vascular lacunae (star) disrupt the organization of muscle bundles and their external connective tissues in a *Krox20-Cre^+/o^*; *Pik3ca*^H1047R/+^ mutant thigh. Bar, 200 μm. (**S**) Disrupted vascular structures (star) were also present in the *Krox20-Cre^+/°^*; *Pik3ca*^H1047R/+^ mutant salivary gland. (**T**) A profusion of small vascular sinuses (star) were enlarged and necrosis was visible (arrow) in the *Krox20-Cre^+/o^*; *Pik3ca*^H1047R/+^ liver. Bar = 100 μm. ao, aorta; la, left atrium; lv, left ventricle; pt, pulmonary trunk; ra, right atrium; rca, right coronary artery; rv, right ventricle, sa, septal artery. Stars, vascular lesions.

*Krox20-Cre^+/o^*; *Pik3ca*^H1047R/+^; *RdT^+/o^* mice, like their *Krox20-Cre^+/o^*; *RdT^+/o^* counterparts, expressed the fluorescent Tomato lineage marker in the smooth muscle of the gonad (**Figure 5**(**H**)); in finely ramified cells, probably reticular fibroblasts, in the spleen (**Figure 5**(**J**)); and in skeletal muscles throughout the body. As expected, cells in the aorta and pulmonary trunk were derived from *Krox20*-expressing progenitors (**Figure 5**(**L**)), but there were also scattered, filamentous cells throughout the walls of the ventricles and to a lesser extent, the atria, corresponding to the sites of developing cardiac vascular lesions in *Pik3ca*^H1047R^ mutant animals (**Figure 5**(**D, I, M**)).

A search for *Egr2* expression in a recent multi-organ database of single adult mouse fibroblasts and vascular wall (mural) cells (https://betsholtzlab.org/Publications/FibroblastMural/database.html) (Muhl et al., 2020) demonstrated that the cells that had expressed *Krox20-Cre* to induce *Pik3ca*^H1047R^ were likely to be endomysial and perimysial fibroblasts in cardiac and skeletal muscle, myelinating Schwann cells and endoneurial fibroblasts in the peripheral nerves, and a previously undefined subtype of vascular pericytes, fibroblasts and smooth muscle. Consistently, fascicles of the peripheral nerves, including autonomic, and a ring of probable pericytes (Topilko, 2019) at the base of hair follicles in the non-glabrous skin strongly expressed RdT in adulthood (**Figure 5**(**L, N, O**)). These lineage observations are compatible with the independent single-cell data (Muhl et al., 2020) and account for the unexpectedly widespread, progressive vascular phenotypes in *Krox20-Cre^+/o^*; *Pik3ca*^H1047R/+^ mutant mice.

Multipotent cells in fibrous connective tissues, characterized by limited or ongoing *Egr2* transcription, appear to mediate hypertrophy of the vasculature and spleen in response to exposure to an as-yet undefined interval of sustained PI3K signaling.

### 3.5 Egr2-driven *Pik3ca*^H1047R/+^ expression induces postnatal pituitary and intramuscular vascular remodeling

Since *Krox20-Cre^+/o^*; *Pik3ca*^H1047R/+^ mice survived to adulthood, we made a cursory examination of the pituitary gland for morphological or functional anomalies to compare to the *Wnt1-Cre^+/o^*; *Pik3ca*^H1047R/+^ phenotype. Lineage tracing with the *RdT* allele as above demonstrated that most if not all cells of the adenohypophysis, unlike the neurohypophysis, had once expressed the Krox20 transcription factor (**Figure 6**(**A-F**)). Examination of the capillary network in control mice with Pecam1 (CD31) immunofluorescence in confocal microscopy showed that some Pecam1+ cells also showed Tomato fluorescence and thus had also once expressed Egr2/Krox20 (**Figure 6**(**C-F**), arrowheads). Intriguingly, Pik3ca-mutant mosaic pituitaries had visibly fewer RdT+ cells and a concomitant decrease in Pecam1+/ RdT+ cell density (**Figure 6**(**G-J**), n=7), although not in organ size. Cavernoma-like lesions were never observed in *Krox20-Cre^+/o^*; *Pik3ca*^H1047R/+^ pituitaries (n=22).

**Figure 6.**
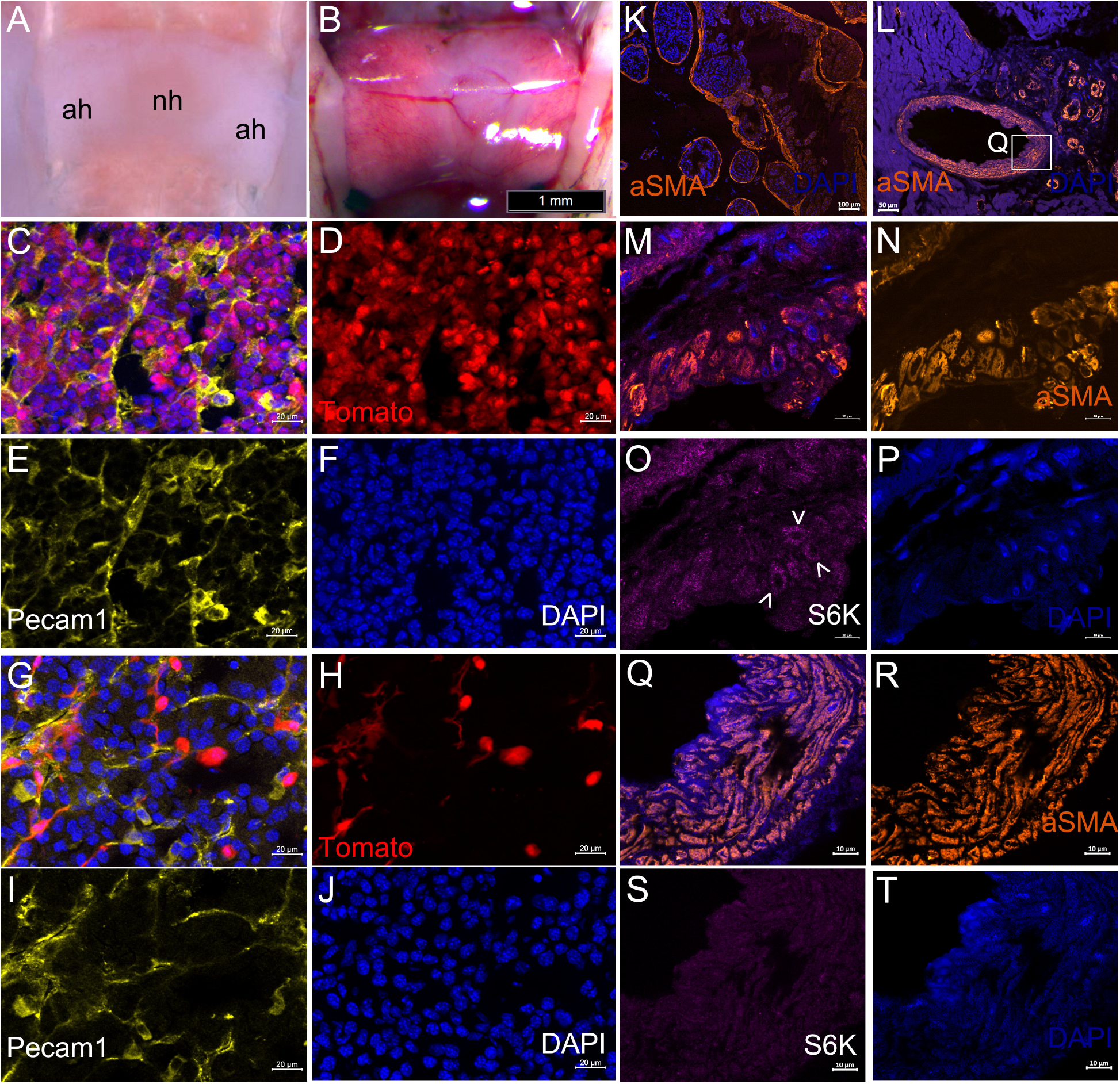
Lineage tracing and immunofluorescence in *Krox20-Cre^+/o^*; *Pik3ca*^H1047R/+^ adult mutant mice. (**A**) Dorsal view of control pituitary after removal of brain and meninges. (**B**) Dorsal view of *Krox20-Cre^+/o^*; *RdT*^+/o^ pituitary during dissection after removal of brain, showing highly fluorescent adenohypophysis under visible light. Bar, 1 mm. (**C-F**) Section through adenohypophysis of *Krox20-Cre*^+/°^; *RdT*^+/o^ mouse, showing normal cell density and that most or all cell types, including perivascular nuclei, had expressed *Krox20*, unlike the sparse recombination observed in the neurohypophysis (not shown). (**C**) Merged (**D**) Tomato (**E**) Pecam1 (CD31) (**F**) DAPI fluorescence. Bar, 20 μm. (**G-J**) Section through adenohypophysis of *Krox20-Cre^+/o^*; *Pik3ca*^H1047R/+^; *RdT^+/o^* mouse, showing reduced cell density accompanied by a striking reduction in cells that had expressed *Krox20*. Bar, 20 μm. (**K, L**) Expression of alpha-smooth muscle actin (aSMA, orange) around vascular tumors (**K**) and in a coronary artery and surrounding telangiecstasias (**L**). Area magnified in (**Q**) indicated. (**M-P**) Mural structure in representative vascular anomaly. Slight increase of phosphorylated S6 kinase (**O**, purple, arrowheads) in a few among the disorganized and unusually shaped cells of the aSMA-expressing vascular wall (**N**, orange). (**M**) Merged. (**P**) DAPI. Bar = 10 μm. (**Q-T**) Mural structure of the coronary artery in (L). No apparent increase of phosphorylated S6 kinase (**S**, purple), but presence of disorganized and unusually shaped cells in the aSMA-expressing vascular wall (**R**, orange), although some laminar structure is still present. (**Q**) Merged. (**T**) DAPI. Bar = 10 μm.

A recent report implicated increased PI3K signaling in the formation of cerebral cavernous malformations (CCMs) and phosphorylated S6 (p-S6) ribosomal protein expression as its endothelial intermediary (Ren et al., 2021). We therefore sought, but did not observe, similarly increased expression of p-S6 in the vascular endothelium of *Krox20-Cre^+/o^*; *Pik3ca*^H1047R/+^ mice (**Figure 6**(**O, S**)). Increased expression was sometimes observed in the abnormally shaped walls of muscular cavernomas, which co-expressed aSMA (**Figure 6**(**M, O**, arrows)). No increase was seen in the thickened, disorganized smooth muscle walls of the coronary artery (**Figure 6**(**Q, S**)).

### 3.6 Melanocytic and other tumors

In many *Krox20-Cre^+/o^*; *Pik3ca*^H1047R/+^ mutants, vascular anomalies were accompanied by widespread, extracutaneous pigmented melanocyte deposits (**Figure 7**(**A**)). In the meninges of the head, although some melanocytosis is physiological in mice (Gudjohnsen et al., 2015), the olfactory lobes (**Figure 7**(**B**)) and trigeminal nerves were covered in a melanocytic mesh. Pigmented melanocytes were also conspicuous in the capillary network of the lower incisor gingiva (**Figure 7**(**C**)), which has not been described to our knowledge as a site for extracutaneous melanocytes.

**Figure 7.**
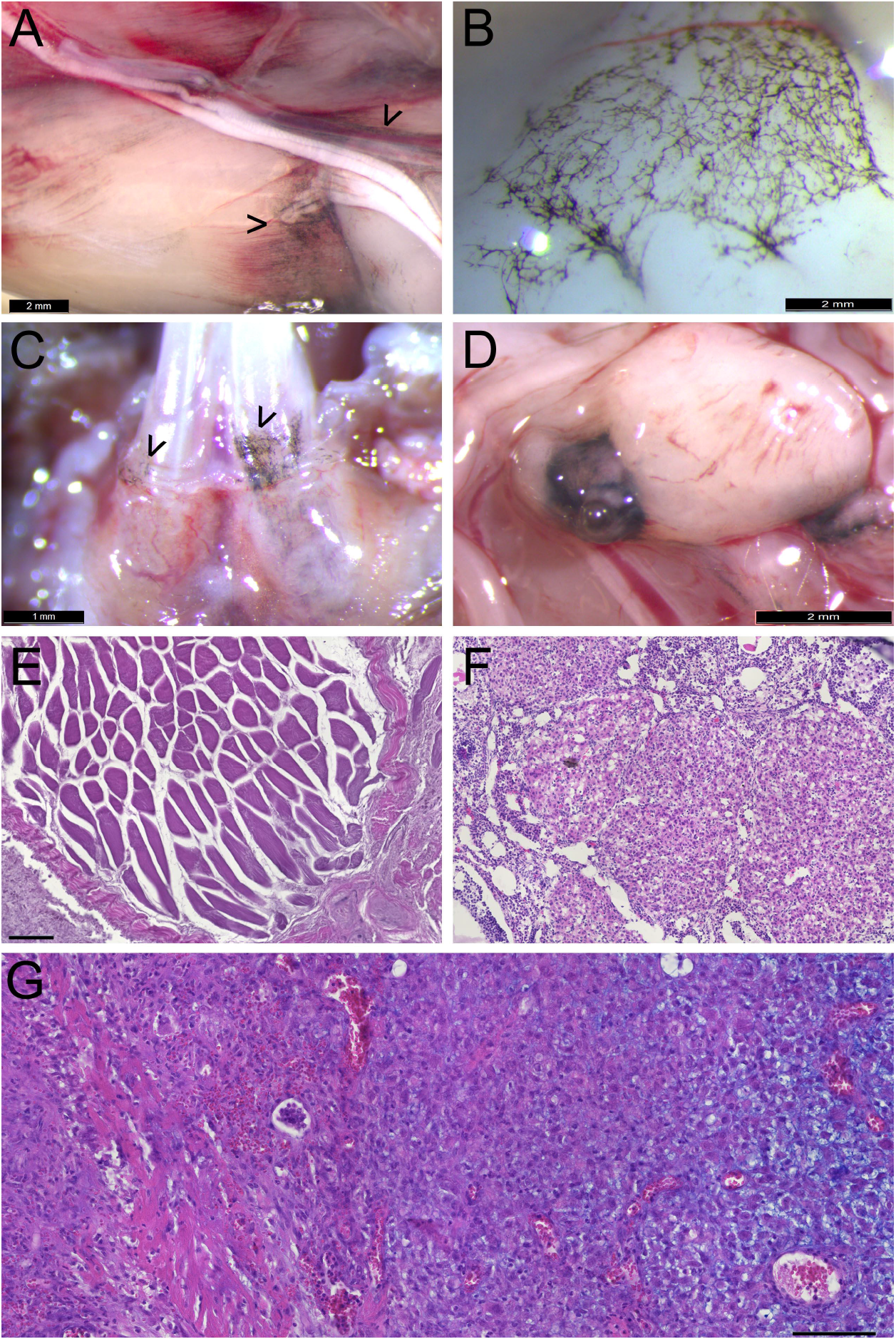
Widespread melanocytic anomalies in conjunction with Krox20- or Sox10-driven expression of constitutively active Pik3ca. (**A**) Extracutaneous pigmented melanocyte deposits along nerves and muscle fascia in *Krox20-Cre^+/o^*; *Pik3ca*^H1047R/+^ mice, close to vascular anomalies. Bar = 2 mm. (**B**) Increased meningeal melanocytosis over ventromedial frontal lobes and olfactory bulbs in *Krox20-Cre^+/o^*; *Pik3ca*^H1047R/+^ mutants, not seen in 4OH-TAM-treated *Sox10-Cre^+/o^*; *Pik3ca*^H1047R/+^ mice. Bar = 2 mm. (**C**) Pigmented, gingival melanocytosis over lower incisors of *Krox20-Cre^+/o^*; *Pik3ca*^H1047R/+^ mutants, not seen in 4OH-TAM-treated *Sox10-Cre^+/o^*; *Pik3ca*^H1047R/+^ mice. Bar = 1 mm. (**D**) Melanoma in *Krox20-Cre^+/o^*; *Pik3ca*^H1047R/+^ mutant mouse near seminal gland. Bar = 2 mm. (**E**) Rhabdomyomatous mesenchymal hamartoma in thigh of *Krox20-Cre^+/o^*; *Pik3ca*^H1047R/+^ mutant mouse. Bar = 100 μm. (**F**) Ovary of *Sox10-CreER^T2^*; *Pik3ca*^H1047R/+^ mouse induced at 15 weeks by 1 mg 4OH-TAM, after 5 days. (**G**) Unpigmented, myxoid melanoma in tail of prenatally induced *Sox10-CreER^T2^*; *Pik3ca*^H1047R/+^ mouse, after nearly one year. Bar = 100 μm.

Four adults developed melanocytic tumors in addition to their vascular anomalies (**Figure 7**(**D**)). These regularly invested distant lymph nodes and were found in multiple sites, but without typical melanoma-like metastasis to brain, liver or lung, where tumors were never observed. A rhabdomyomatous mesenchymal hamartoma was also observed in the inner thigh of one mutant mouse (**Figure 7**(**E**)).

Peripheral nerve Schwann cells must express Krox20 to myelinate, and can shuttle between a differentiated, myelinating mature and a non-myelinating, immature state depending on tissue needs (Decker et al., 2006). Furthermore, immature Schwann cells give rise to melanocytes under both physiological or pathological conditions (Adameyko et al., 2009). We hypothesized that Schwann cells that had expressed Krox20 and undergone a phenotypic switch after expression of constitutively active Pik3ca in mutants could be the source of the widely distributed extracutaneous melanocytes. To test this *in vivo*, but bypass the lethal phenotype of *Wnt1-Cre*; *Pik3ca*^fl(H1047R)/+^ mice, we targeted nerve-resident peripheral glial precursors (Deal et al., 2021) by crossing *Sox10-CreER^T2^* mice with floxed *Pik3ca*^fl(H1047R)/+^ mice. This allowed us to induce *Pik3ca*^fl(H1047R)^ expression at will after exposing *Sox10*-expressing cells to the metabolically active tamoxifen derivative, 4-hydroxy-tamoxifen (4OH-TAM).

Nine *Sox10-CreER^T2^*; *Pik3ca*^H1047R/+^ mice (eight female, one male) from four litters were injected with 1 mg 4OH-TAM at 15-19 weeks and compared to eight similarly treated *Sox10-CreER^T2^*; *RdT* mice (five female, three male) and one female *Pik3ca*^fl(H1047R)/+^ controls of the same age. Within five days, two mutants had died and three others had attained a humane endpoint and were euthanized. The remaining mice appeared healthy at 14 days post injection. Gross examination of the *Sox10-CreER^T2^*; *RdT* mice under a fluorescence binocular dissecting microscope demonstrated effective recombination had been induced in all. Autopsy did not reveal the causes of morbidity or mortality. The superficially pigmented axillary and Peyer’s patch lymph nodes had enlarged germinal centers, and the ovaries were cystic in one deceased and all three surviving female *Pik3ca*-mutant mice (**Figure 7**(**F**)). No widespread melanocytosis or vascular tumors were noted.

Although unlikely to be a cause of death, we conjectured that constitutively active PI3K signaling in post-migratory NC and other *Sox10*-expressing cells may predispose the later Schwann cell progenitors to melanocytosis. To test this idea, *Sox10-CreER^T2^* males were mated to three *Pik3ca*^flox(1047R)/+^ females and the pregnant dams treated at E18.5 with 4OH-TAM. This led to recovery of a total of ten live births: five *Sox10-CreER^T2^*; *Pik3ca*^H1047R/+^ (three female, two male), four *Sox10-CreER^T2^* and one male *Pik3ca*^H1047R/+^ mouse. In contrast to the induction of *Pik3ca* mutation in adulthood, these prenatally induced animals were followed without incident for up to 1 year, when one male mutant rapidly developed an unpigmented, circumscribed tail tumor of 5 mm in diameter and showed signs of distress. After euthanasia, varied cellular elements including smooth muscle and mucin-containing myxoid zones that stained with Alcian blue were found in the tumor (**Figure 7**(**G**)).

In all of these models, expression of the Pik3ca(H1047R) allele during development appears to promote hyperplasia and tumor formation in cell lineages competent to give rise to connective and support tissues. Within post-migratory Schwann cell progenitors expressing Krox20 and Sox10, the proliferative state also promotes melanocytosis.

## 4 Discussion

The work presented above shows some of the complex developmental effects of constitutive PI3K signaling on interdependent organ systems in the context of mosaicism. These range from the jaws to the brain and from blood vessels to pigment cells. Three new, mechanistically related mouse models complement one another to demonstrate pathogenic diversity.

### 4.1 Tumor-like vascular malformations arise from mutant perivascular cells

*PIK3CA* gain-of-function mutations lead to constitutive activation of AKT downstream of the TEK angiopoietin-1 receptor in human vascular endothelial cells (Limaye et al., 2015; Kobialka et al., 2022) as well as in endothelial cells of targeted mouse models (Castillo et al., 2016). Nearly half of sporadic venous malformations (VMs) bear activating *TEK* mutations, while most others express activating *PIK3CA* H1047R, or E452K or C420R mutations, in a mutually exclusive manner (Castel et al., 2016). These, particularly H1047R, are also the most frequent hotspot mutations for malignancies in nearly 50 sites and in over 21,000 samples curated by COSMIC (v95) to date (Tate et al., 2019; Avramović et al., 2021).

The work we present here is the first to demonstrate that activation of a complex signaling pathway by the identical mutation in abluminal pericytes, fibroblasts and vascular smooth muscle, rather than endothelial cells, also can induce congenital vascular malformations and tumorigenesis. Paracrine signaling between endothelial and mural cells is necessary for tissue-appropriate, functional blood and lymphatic vessel assembly. For example, pharmacological PI3K inhibition rescues inducible arteriovenous malformations in the context of an inducible animal model for a recurrent transforming growth factor-b (TGF-b)/bone morphogenetic protein (BMP) signaling pathway gene mutation, known to cause hereditary hemorrhagic telangiecstasia (Ola et al., 2016).

p-S6, a downstream effector of AKT/mTOR, was no longer active in most of the adult cavernoma-like vascular lesions we observed in mice with mutant Pik3ca in cells having once expressed Egr2/Krox20. This contrasts with findings from a recent mouse model for CCMs due to *Pik3ca* mutation (Ren et al., 2021). A punctual, rather than sustained, earlier stimulus may be sufficient to launch some processes of tissue overgrowth; p-S6 is only activated during the process of angiogenesis in the presence of exogenous growth factors in *Pik3ca*-mutant retinal endothelial cells (Kobialka et al., 2022). Activated MAPK effectors may also participate in vascular pathogenesis, as seen in a case report of a child with Noonan syndrome and a retinal CCM (Lallau et al., 2022). Sustained activation of MAPK and PI3K signaling pathways is subject to cell- and tissue-dependent paradoxical repression through compensatory feedback mechanisms, such as upregulation of phosphatases or alternative kinases (Mendoza et al., 2011; Jiang et al., 2022). Further study of perinatal retinal vascularization in our new mouse models may enable definition of the relevant time window, specific downstream mediators and intracellular effectors of these endothelial-mural paracrine exchanges.

The fact that both *Wnt1-Cre*; *Pik3ca*^H1047R/+^ and *Krox20-Cre*; *Pik3ca*^H1047R/+^ mice both present vascular anomalies within cardiac tissue implies that signaling through the PI3K pathway to a cell of NC origin in the heart has an impact on its subsequent paracrine activity, with a similar effect on intracardiac vascular development as in the head and neck. Single-nucleus RNA sequencing of cardiac ventricles and craniofacial mesenchyme in these mouse crosses is a potential approach to identify such vasculogenic factors mediated by lineage-traced, non-endothelial populations.

*Pik3ca*-mutant, lineage-traced NC cells were found in malformed dermal blood vessels over the parietal bone. This observation was remarkable since the NC-derived dermis, vascular pericytes, craniofacial bones and forebrain meninges usually occupy the same territories of the ventral head, face and neck (Etchevers et al., 1999) and share clonal origins in mice (Kaucka et al., 2016). Given that the mouse parietal bone is of mesodermal rather than NC origin, as shown by lineage tracing in a cross of the same *Wnt1-Cre* line with a conditional reporter allele (Jiang et al., 2002b), overgrowth promoted by PI3K signaling may enable some mesenchymal NC derivatives to expand into ectopic cranial regions before birth.

### 4.2 Cell-autonomous PI3K activity affects mesenchymal expansion in the skull

The constitutive activation of PI3K in cells having expressed Wnt1 in the neuroepithelium before NC migration also led to apparently autonomous effects, with macroscopic megalencephaly of the forebrain and midbrain. Of cephalic NC derivatives, only the mesectodermal subpopulation appeared affected (Le Lievre and Le Douarin, 1975). Lineage tracing with a floxed Rosa-tomato fluorescent reporter allele did not show any differences in initial NC distribution after migration into the face and head at E9.5, implicating PI3K signaling in the later differentiation of some cephalic NC-derived mesenchyme into perivascular cells and other connective tissues (Etchevers et al., 2001; Deal et al., 2021). There were no apparent morphological effects at the level of the trunk or in the size of the dorsal root and cranial ganglia in *Wnt1-Cre*; *Pik3ca*^H1047R/+^ neonates, implying that a sustained mesenchymal state predisposes to cell-autonomous effects.

When cranial NC expresses constitutively active Pik3ca, vascular and jaw hyperplasia result. A clear association between vascular and craniofacial overgrowth has been reported clinically for decades, well before molecular genetics caught up (Krings et al., 2007). Facial capillary malformations found in Sturge-Weber syndrome are regularly associated with segmental overgrowth of the orbit or the jaw. This disease is usually caused by constitutively activating, somatic *GNAQ* mutations (Shirley et al., 2013) but rarer somatic mutations, including a pathogenic one in *PIK3CA* (Lian et al., 2014), have been reported. G-protein and PI3K signaling cascades are thus both likely to mediate the exchanges of vascular cross-talk with its surrounding mesenchyme in these evolving skull anomalies.

This excess of NC-derived cranial mesenchyme produced under the influence of oncogenic *Pik3ca* yields both hyperplasic frontal bones and parietal bone hypocalcification. Genetically induced increases in neural crest mesectoderm relative to mesodermal mesenchyme, achieved by tweaking components of the Hedgehog and fibroblast growth factor signaling pathways, also lead to proportionately larger frontal than parietal bones, to the extent of parietal aplasia, as well as palate malformations (Tabler et al., 2016). Differences in PI3K signaling within the osteogenic NC of different vertebrate species and their paracrine influence on the neighboring, ancestral osteogenic mesoderm may be responsible for the diverse morphological specificities of homologous skull bones. These homologies have been a long-standing source of controversy in evo-devo (Noden and Trainor, 2005; Carroll, 2008).

### 4.3 Congenital overgrowth usually does not lead to malignancy but presents its own problems

While nearly half of overgrowth PRD patients in one cohort presented congenital vascular malformations, tissue overgrowth in the vast majority continued to evolve postnatally (Keppler-Noreuil et al., 2014). Somatic mutation of codon H1047 is frequently but not exclusively implicated in asymmetric, multi-systemic *PIK3CA*-related overgrowth disorders such as CLOVES [OMIM 612918; congenital lipomatous overgrowth, vascular malformations, epidermal nevi and skeletal abnormalities] and endophenotypic segmental overgrowth syndromes affecting muscle and fat, or fibroadipose hyperplasia. Another class of PRDs involve megalencephaly with various other features affecting musculoskeletal, vascular, connective and adipose tissues, of which megalencephaly-capillary malformation-polymicrogyria syndrome (MCAP) is emblematic (Lee et al., 2012; Kingsmore et al., 2013; Alcantara et al., 2017). Some patients develop supernumerary, hypertrophic muscles in the upper limbs; these are occasionally bilateral, indicating that the original somatic mutation may have developed in a cell whose progeny entered the paraxial mesoderm cell during gastrulation (Frisk et al., 2019). PI3K/Akt inhibitors have been very promising in clinical trials for these conditions (Dill et al., 2014; Roy et al., 2015; Venot et al., 2018; Hori et al., 2020; Kobialka et al., 2022)

Malignant tumors in PRD patients are surprisingly rare, given that they express proven oncogenic mutations, and this finding was borne out in our mouse models. However, the resultant malformations themselves were not always compatible with viability.

We have observed that benign vascular and/or hamartoma-like tumors arise postnatally in mice expressing constitutively active Pik3ca in a mosaic fashion, in cells having expressed the *Krox20/Egr2* or *Sox10* transcription factors. Both of these transcription factors are hallmarks of so-called “Schwann cell precursors” or SCP. SCP are NC-derived, non-myelinating cells that reside along or at the terminal ends of peripheral nerves, and that can respond to environmental changes such as injury or inflammation by differentiating into myelinating Schwann cells and endoneurial fibroblasts but also melanocytes in rodents (Adameyko et al., 2009). Interestingly, these resident, poised cells also normally contribute extracutaneous melanocytes to the heart, inner ear, ocular choroid plexus, to the perimysial layers of some skeletal elements (unpublished observations) and the CNS meninges (Kaucka et al., 2021).

Even without additional PI3K activity, modest melanocytosis was seen in the hypothalamic meninges and membranes surrounding the trigeminal nerves of adult pigmented *Krox20-Cre;RdT* mice. In contrast, agouti or black tamoxifen-induced *Sox10-CreER^T2^* control mice had only normal meningeal pigmentation (Gudjohnsen et al., 2015). Dosage reduction of Egr2 may thus be a prerequisite to SCP plasticity, as the *Krox20-Cre* allele replaces one copy of *Egr2* with *Cre* (Voiculescu et al., 2000). Neither early nor late induction of constitutively active PI3K signaling in *Sox10-CreER^T2^*; *Pik3ca*^H1047R/+^ mice leads to melanocytosis, but rather to other tumors. Our observations support other recent studies that highlight the importance of positional and lineage context for the neoplastic potential of oncogenic mutations (Baggiolini et al., 2021) extending beyond MAP kinases to the PI3K signaling pathway.

The partial commitment in multipotent progenitors at any stage of life, rather than their original lineage, seems to make them particularly vulnerable to overgrowth as an effect of constitutively active PI3K signaling.

### 4.4 Pik3ca-related disorders may encompass previously unsuspected pathologies

The wide variety and range in severity of PRD phenotypes is attributed in part to the location of the cells bearing the mutation and to the proportion of cells affected in each of any given patient’s tissues. We identified effects of constitutive PI3K signaling on pituitary and palatal development that are not features of the diverse PRDs already identified to date.

Adrenal insufficiency could contribute to perinatal mortality in our *Wnt1-Cre*; *Pik3ca*^H1047R/+^ mice, since when constitutive Akt signaling is induced in the embryonic ectoderm, crucial proteins for differentiation of the corticotropic cell lineage, such as Bmp4 and Tbx19, are significantly down-regulated in mice (Segrelles et al., 2008). If so, this would be another measure of paracrine NC effects on pituitary development. The rapid death of a subgroup of *Sox10-CreERT2*; *Pik3ca*^H1047R/+^ mice after 4OH-TAM induction may also be due to adrenal crisis. The nature of the paracrine activity exerted by NC-derived cells, and its effects on adrenal function, could be further investigated in these mouse models.

Lineage-traced *Krox20-Cre; RdT* mice showed a much broader distribution of cells in the body that had once or continued to express *Egr2* than previously described. Evidence exists that some of these are cutaneous vascular pericytes derived from Schwann cell precursors (SCP), themselves from a NC-derived “boundary cap cell” population residing adjacent to the spinal cord (Gresset et al., 2015; Topilko, 2019). However, many also appear to be specialized fibroblasts primed by this transcription factor (Muhl et al., 2020). The cutaneous SCP-derived pericyte may in fact be a specialized myofibroblast, a subject for future studies in fibroblast diversification.

The sites of predilection for *Krox20-Cre*; *Pik3ca*^H1047R/+^ vascular lesions were within the *panniculus carnosus* and epaxial skeletal muscle groups, compatible with intramuscular hemangioma, a tumor rather than a vascular malformation (Tan et al., 2007; Kurek et al., 2012). Unlike soft-tissue angiomatosis, we did not observe a mature adipose or lipomatous component to these lesions, or indeed in any of the other described mouse models. However, adipocytes do collect around peripheral nerves (**Figure 5(A-B)**) in our *Krox20-Cre*; *Pik3ca*^H1047R/+^ mice. This observation may be relevant for the study of fibrolipomatous hamartoma, which like other PRDs is associated with overgrowth of the innervated territory (Marek et al., 2021). It also helps explain why mutations in *PTEN* are frequently associated with intramuscular vascular anomalies, since they lead indirectly to PI3K pathway activation (Tan et al., 2007; Ho et al., 2012).

Recently, gain-of-function *Pik3ca* mutations have been shown to be sufficient to drive small, postnatal capillary hemangiomas in brain endothelial cells and are necessary, in combination with mutations of known CCM genes in mice or in humans, for the development of large postnatal cavernomas (Hong et al., 2021; Ren et al., 2021). Should *PIK3CA* mutations also be confirmed in human intramuscular hemangiomas or fibrolipomatous hamartomas, they would be the functional and tumoral counterpart of soft-tissue hamartomas due to *PTEN* mutations (Kurek et al., 2012; Luks et al., 2015; Tachibana et al., 2018). In these congenital and predisposing conditions, evolving anomalies are not always surgically accessible and can be aggravated by incomplete resection. Targeted inhibitors of distinct pathway levels, locally infused and/or in combination, show great promise and can now be tested in a wider range of tailored animal models.

## 7 Author Contributions

EM, MM and ML planned and performed mouse crosses and dissections, and undertook the histology and immunofluorescence experiments as well as microscopy. AP and KH acquired and analyzed the data for Figure 2. GM and AB provided reagents and expertise on PI3K signaling. TF dissected pituitary glands and aided in the interpretation of the pituitary sections. NM reviewed the histology. HCE contributed the *in situ* hybridization and microscopy, supervised statistical analyses with EM, obtained funding and wrote the manuscript. All authors reviewed the final manuscript.

## 9 Acknowledgments

The authors acknowledge all the members of the DIP-NET team for ensuring continuity under challenging circumstances. They also thank Adeline Ghata and the rest of the MMG animal platform for their precious assistance, Drs. Piotr Topilko and Stéphane Zaffran for the *Krox20-Cre* mouse line, and Dr. Lotfi Slimani for help generating the micro-CT data at the PIV core imaging facility (http://piv.parisdescartes.fr/).

## 10 Contribution to the field statement

In this paper, we have developed multiple new mouse models to test the developmental function of a gene whose mutations are frequent and well known to cancer researchers, called *PIK3CA.* The H1047R mutation is present in 4 out of 10 common malignancies due to this gene, permanently activating the enzyme that it encodes and driving aggressive tumor growth. We have carefully observed and described the anatomical and molecular characteristics of many new malformations that can also be caused by the same oncogenic mutation. Mutations of *PIK3CA* have also been identified over the last decade in numerous rare disease syndromes with overlapping symptoms, among which musculoskeletal, brain and vascular malformations are regularly observed. By restricting PIK3CA activity to specific subsets of cells in the mouse, it was possible to assess their changed abilities to make or influence other cell types. Mutant loose (mesenchymal) connective cells are most vulnerable to causing changes in tissue shape and size, or to developing cancer. These mouse models indicate additional candidate diseases in humans where PIK3CA may be locally active, opening new potential applications for existing treatments.

